# Real-time spike sorting with 3D neural probe and triangulation localization

**DOI:** 10.1101/2025.03.30.645752

**Authors:** An-Cheng Yang, Kuan-Peng Chen, Jia-Hao Zhang, Kuo-Hsing Kao, Wen-Jay Lee, Myles McLaughlin, Nan-Yow Chen, Jyh-Jang Sun

**Author notes:** Equal contribution (An-Cheng Yang, Kuan-Peng Chen).

## Abstract

A key challenge in correlating neuronal activity with brain function is the limited sampling probability of neuronal activity in real time. It is crucial to increase the sampling probability substantially and in real time. We hypothesized that 3-dimensional (3D) neural probes offer faster and stronger prospects for cell yield than 2D electrode arrays. We simulated a 1000-neuron neuronal network, mimicking the granular layer of the barrel cortical column, recorded signals from inserted 384 electrodes (organized in 3D or 2D), and sorted units using Kilosrt or triangulation localization. We demonstrated that 3D electrode arrays converge more space for triangulation spike sorting than 2D probes do. 3D neural probes, together with triangulation, could isolate up to 80% of the simulated 1000 neurons (as ground truth) and have a cell yield of up to 5, which is, to the best of our knowledge, significantly higher than standard 2D electrodes with Kilosort or triangulation. With a signal-to-noise ratio (SNR) of 10, which is close to the real world, the simulation data suggest that 3D electrode arrays in a face-centric cubic (FCC) arrangement provide a better cell yield. However, larger background noise (e.g. an SNR of 1, which can be improved with lower electrode impedance) has a stronger impact on the triangulation spike sorting. Since only the peak value of spikes are required for triangulation localization, the computing loading is much less than spike waveform-based spike sorting approach. Thus, combining 3D electrode arrays with triangulation localization is ideal for real-time spike sorting. Thus, we demonstrated that adding one more dimension in designing neural probes can dramatically increase cell yield and speed up isolating neuronal unit activity. We, for the first time, provide a tool for utilizing computer simulations to optimize the design of neural electrode arrays before time-consuming probe fabrication.

## Introduction

A key challenge in correlating neuronal activity with brain function is the very limited sampling probability of neuronal activity, i.e., typically 100 out of 15,000 cells in a cortical column. Thus, it is crucial to substantially increase the sampling probability. Extrapolating from existing published work in terms of cell yield of various neural probes[1–3], we hypothesize that dense 3D neural probes offer strong prospects for this.

When neural signals are acquired by neural probes, i.e., microelectrode[4], stereotrode[5], tetrodes[6,7], and silicon probe[8], neuroscientists start a procedure to classify which recorded neuronal spikes correspond to which neurons, called spike sorting[9], for review papers see[10,11]. The typical approach to spike sorting consists of an initial spike detection, extracting features of spike waveforms, and finally, a clustering analysis on those features is performed. After the clustering, the quality of each individual cluster is assessed by a human operator, commonly using phy[12]. Nowadays, spike sorting is routinely used in many neuroscience labs and several toolboxes, i.e. Klustakwik[13], Klusta[12], Kilosort[14], MountainSort[15], SpyKING CIRCUS[15,16], have been developed to speed up and simplify this procedure. However, the problems in the procedure of spike sorting, i.e., identifying bursting cells, clarifying overlapping of spike waveforms, merging/splitting clusters, contamination from unknown spikes, and the time spent on manual curation, remain unsolved. Moreover, after spike sorting many of the recorded spikes (approximately 50%) are discarded, due to the poor separation of spike waveform features. Therefore, we need to collect more spike waveform features in neural circuits to significantly increase the yield of spike sorting. Although high-channel-count probes, i.e., Neuropixels probe[2] and NeuroSeeker probe[17], have been developed (up to 384 and 1344 recording sites), those problems remain unsolved.

The key for spike sorting relies on the assumption that different cells exhibit spikes that have unique amplitudes and waveforms. Therefore, multiple electrodes distributed in 3D space could record better separable spike features, i.e., tetrode[1]and 3D silicon probe[18–21]. Although 3D silicon probes by stacking 2D probes have been developed[18–20] and commercialized (Neuronexus, ATLAS Neuroengineering), they cause severe tissue damage and are difficult to insert into the brain and therefore unfortunately are limited in their usage in larger brain regions. Recently, neural probes made from polyimide or SU-8 into 3D-shape have been developed, i.e., cylindrical probes[22,23], folded probes[24–26], self-deployable probes[27]^,[28]^, for minimizing tissue damage[29,30] and extensions of the recording sites. To this end, unfortunately, no dense 3D polyimide neural probes with great coverage have been invented.

It is impossible to fully understand how the brain operates and interacts at the level of the individual neuron in naturally behaving animals unless we first solve the problems of spike sorting and invent a novel tool for the brain-computer interface at the cellular level. Thus, we need a simple, fast, and accurate spike sorting approach for closed-loop experiments to fine-tune neural networks in real-time.

Here, we first ask whether electrodes organized in 3D can sample significantly more well-isolated neurons than in 2D by computer simulations. We further ask how electrodes organized in 3D give us the best performance in terms of cell yield. Due to the particular structure of 3D electrode arrays, we evaluated the performance of various electrode arrays not only using Kilosort[14] but also utilizing triangulation spike sorting, based on positioning neurons (namely triangulation spike sorting). The triangulation spike sorting is based on each neuron’s unique location relative to recording electrodes through a method of triangulation. This is most effective if signals are sampled along orthogonal axes, requiring a 3D electrode constellation. Theoretically, spike sorting by 3D triangulation would greatly speed up the procedure of spike sorting. Previous reports on spike sorting by triangulation have been limited to planar 2D neural probes[31] or tetrodes[32]. Dense and structured 3D neural probes have not been explored yet, due to the lack of suitable fabrication technology. Even though a suitable 3D neural probe is expected to be able to map most neurons in the volume covered by its electrodes, e.g. having 4 electrodes in 3 axes in a cube with an edge length of 50 μm should provide recordings from 8-12 neurons. Based on this argument, we expect to isolate several hundreds of cells with a 126-252 electrode 3D neural probe. Through the use of optimized algorithms, we expect that triangulation spike sorting will become fast enough to perform closed-loop experiments at the cellular level in real-time. Taken together, such closed-loop experiments, where causal and conclusions related to the role of neuronal activity with behavior performance can be drawn. This novel tool is anticipated to dramatically accelerate the pace at which neuroscientific breakthroughs take place and will eventually lead to new clinical applications and treatments.

## Results

### 3D electrode arrays converge more space for triangulation spike sorting than 2D probes do

Our overall objective is to ask whether 3D neural probes can outperform cutting-edge 2D neural probes, i.e., Neuropicels probes (NPX). To accomplish this, we started from simulations of neural networks to determine electrode layouts and arrangements in 3D where we can obtain the best yield of isolated neurons and compare the results of NPX. To benchmark the performance of steric neural probes on isolating single unit activities, we first adopted the following performance index: exploration volume of electrodes (light blue in Fig. 1A) and significant volume of electrodes (darker blue in Fig. 1A). We assumed that significant volume and the ability in triangulation of steric probes is highly correlated, so we adopted significant volume as one of the performance indexes in this study. Exploration volume is the union of the sensing volume of all electrodes in space. Significant volume is the intersection of sensing volume of at least 3 electrodes in space. Second, we further compared the significant volume of various steric probe patterns, e.g., simple cubic, faced-centric cubic, etc., (Fig. 1B), with various electrode pitches, i.e., 40 to 150 um, and various electrode sensing radii, i.e., 75 to 150 um, based on current fabrication technology. Finally, we gave suggestions in the design of 3D probes for obtaining more significant volume.

**Figure 1.**
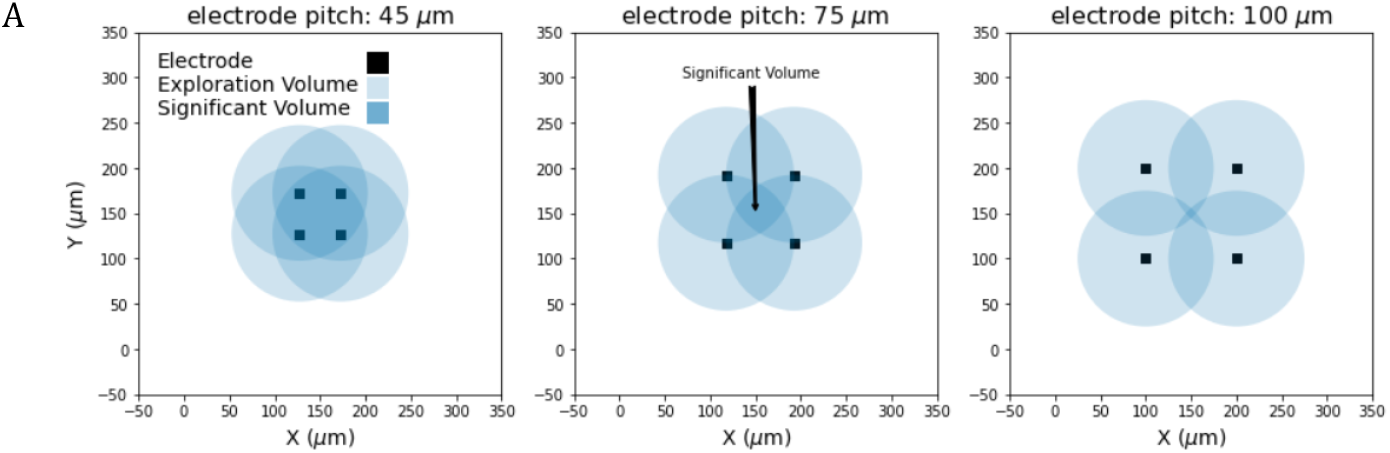
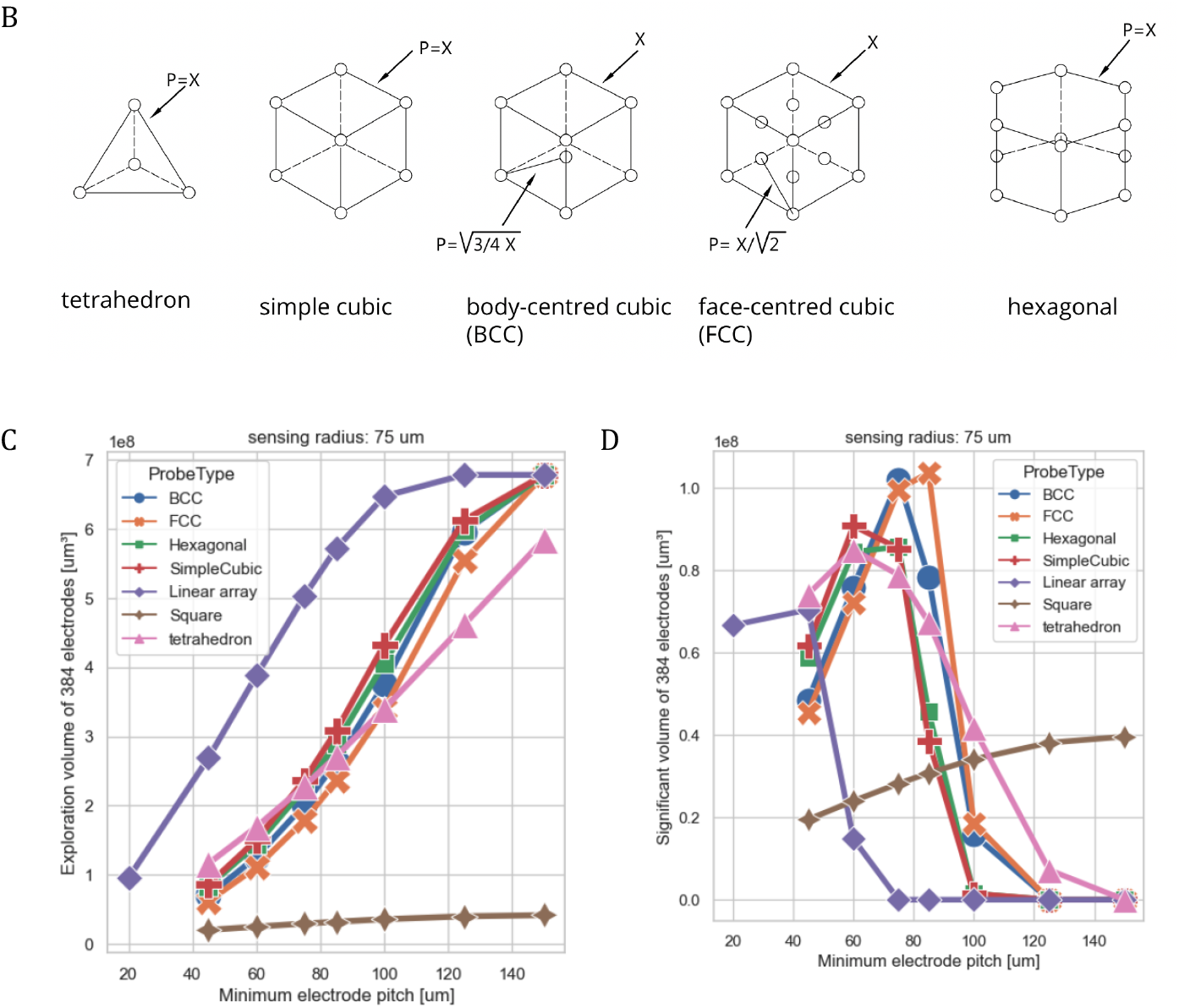
Illustration of exploration volume and significant volume in various 3D electrode arrays. (A) Illustration of the concept of exploration volume and significant volume with a simple cubic electrode array. For better illustration, the electrode array in 3D is projected to 2D. Black dots indicate electrodes, and black squares indicate electrodes, with their sensing radius of signals, i.e. 75 μm, in light blue. Union of light blue area and darker blue is defined as exploration volume. Intersection area with at least from 4 depicted in dark blue, which is defined as significant volume. Exporation volume increases while the electrode pitch increases, i.e. 45 μm to 100 μm. But significant volume decreases while the electrode pitch increases, i.e. 100 μm to 45 μm. (B) 5 types of 3D structures are proposed, i.e. simple cubic (SC), body-centered cubic (BCC), face-centered cubic (FCC), hexagonal, and tetrahedron. (C) Exploration volume increases in all types of 3D electrode arrays. Note that the linear array (Neuropixels-like) reaches peaks with electrode pitch of 100 μm. (D) Significant volume of all 3D electrode arrays dramatically increases and reaches peaks at electrode pitch of 85-100 um but decreases afterwards. Note that the linear array (Neuropixels-like) reaches the peak at electrode pitch of 45 μm.

The neural probe features hundreds of recording electrodes that can individually acquire AP (action potential) and LFP (local field potential) signals from nearby neurons. High dense recording sites of commercial probes(e.g., NPX) yield a huge amount of spiking activity of neurons, but the planar silicon-based shank blocks the signals coming from those neurons behind the probe. The exploration volume of a planar probe, then, in theory, has shrunk by almost half. A steric probe with double-side electrodes expands the exploration volume in three-dimensional space. We use “voxel” representation in 3D computer graphics to calculate the volume of the explorable space of a probe. We adopted 1μm as the size of voxels. Based on the observation in previous electrophysiological experiments, we assumed an average of 75μm to 150μm of sensing radius in signal detection of electrodes, see Fig. 1CD. We defined the overall voxels where the overlapping degree is greater than 3 as significant volume.

We further introduced a concept from material physics that atoms could be self-organized into particular 3D structures, i.e. simple cubic (SC), body-centered cubic (BCC), face-centered cubic (FCC), hexagonal, and tetrahedron, as well as NPX-like 2D electrode array (called linear) and MEA-chip-like 2D electrode array (called square, similar to MEA chip with a 6×6 electrode array in 2D, MCS, Reutlingen, Germany). We aim to test whether electrode arrays in 3D outperform electrode arrays in 2D in terms of triangulation spike sorting. We then computed the exploration volume and significant volume over different electrode pitches (i.e. 45, 60, 75, 85, 100, 125 and 150 μm) with 75μm of electrode sensing radius (Fig. 1CD). Note that the electrode pitch of NPX-like (linear) started from 20μm, as defined in NPX 1.0[2]. As expected, the exploration volume increases in all types of 3D electrode arrays (Fig. 1C) with the increases of electrode pitch. On the other hand, significant volume of all 3D electrode arrays dramatically increased and reached peaks at electrode pitch of 85-100 μm but decreased afterwards (Fig. 1D). Not that the significant volume of the linear array (Neuropixels-like) reaches peaks when the electrode pitch is 45 μm and then decreased afterwards. FCC and BCC with electrode pitch of 90 μm and 75 μm gained the largest significant volume among all types of 2D and 3D electrode arrays. Hexagonal, simple cubic, and tetrahedron reached their peaks at electrode pitches of 85 μm. Note that the linear array (Neuropixels-like) and square gains a relatively smaller significant volume than the others.

These simulated data suggested that steric probes provide one more degree of freedom than planar probes (i.e. linear array and square) do, which can endow researchers with better spatial features for spike sorting, e.g. triangulation spike sorting.

We further asked whether a larger sensing radius (i.e. 100-150 μm) of electrode (with lower electrode impedance) can give us larger significant volume for triangulation spike sorting and ask which types of 3D electrode arrays benefit from it most, and with which electrode pitches. As expected that the larger sensing radius is, the larger significant volume is obtained (Fig. 2). We found that the significant volume increased when the sensing radius increased to 100, 125 and 150 μm. Similar to the result of 75-μm sensing distance,FCC and BCC gained the largest significant volume. FCC and BCC with electrode pitch of 100, 125, and 150 μm gained the largest significant volume, corresponding to sensing radius of 100, 125, and 150 μm. Thus, in theory, if the sensing radius of electrodes increases from 75 μm to 150 μm one can obtain ca. 8 times more significant volume, meaning a possibility of isolating ca. 8 times more neurons.

**Figure 2.**
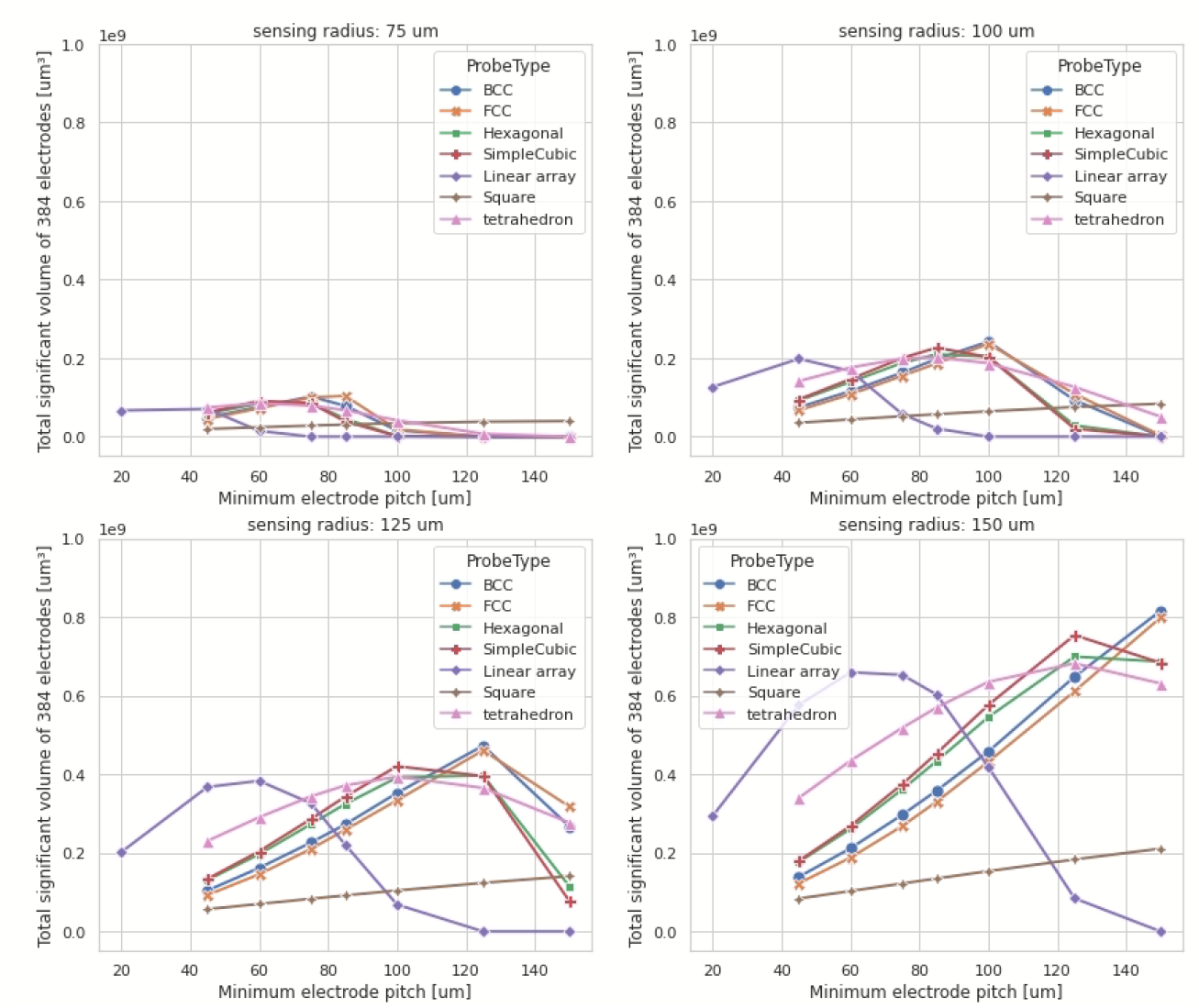
Illustration of significant volume of various 3D types electrode arrays with electrode sensing distance of 75 μm (as shown in Fig. 1D but rescaled here for better comparison to others), 100 μm, 125 μm and 150 μm. Significant volume of all 3D electrode arrays dramatically increases while sensing radius increases. FCC and BCC with electrode pitch of 100, 125, and 150 μm gain the largest significant volume, corresponding to sensing radius of 100, 125, and 150 μm.

### 3D electrode array outperforms 2D probes in isolating neurons

In order to further investigate whether electrodes distributed in 3D, i.e. FCC or BCC (Figs. 1 and 2, with higher significant volume), can isolate a substantially higher number of neurons than planar probes, i.e., Neuropixels probe (with less significant volume). We addressed this question firstly by constructing simulated neural networks, a cube of 646μm×646μm×3970μm with 1000 neuron cells in it to mimic neural networks in a cortical barrel column as a ground truth (See Materials and Methods and Fig. 3A). Secondly, we “insert” one of the 5 predefined 384-channel 3D electrode arrays (i.e. SC, FCC, BCC, Hexagonal, and Tetrahedron) and 2 types of 384-channel 2D electrode arrays (i.e. linear array and square), see Fig. 3B, and then “record” the simulated field potentials, including spike activities, with sample rate of 30,000 samples per second (see Fig. 3C). Additional, in order to mimic more realistic ephys recordings, we inject different levels of white noise (i.e. signal-to-noise ratio of 1, 2, 5, 10, 20, and 40, as well as no injected raw data) to the recorded signals. Finally, we isolated spiking units using custom modified Kilosort 2.0 for 3D electrode arrays data (named “Kilosort 2.0-3D”, see Materials and Methods) and triangulation spike sorting (see Materials and Methods). Out goal is to benchmark their performance on isolating single unit activities among the 5 predefined 3D electrode arrays and 2 types of 2D electrode arrays. Since Kilosort 2 was designed for 2D electrode arrays, we added one more dimension and validated its accuracy for 3D electrode arrays with different rotating angles in 3D (see Fig. S1). The second proposed spike sorting approach proposed here is based on the method of triangulation, making use of the fact that each neuron has a unique location relative to the recording electrodes[33]. In this study, we further apply the triangulation spike sorting from 2D-electrode-arrays spiking data to 3D-electrode-arrays spiking data. This approach is most effective if the signals are sampled along orthogonal axes in 3D, which is the principal reason why we pursue a 3D electrode constellation.

**Figure 3.**
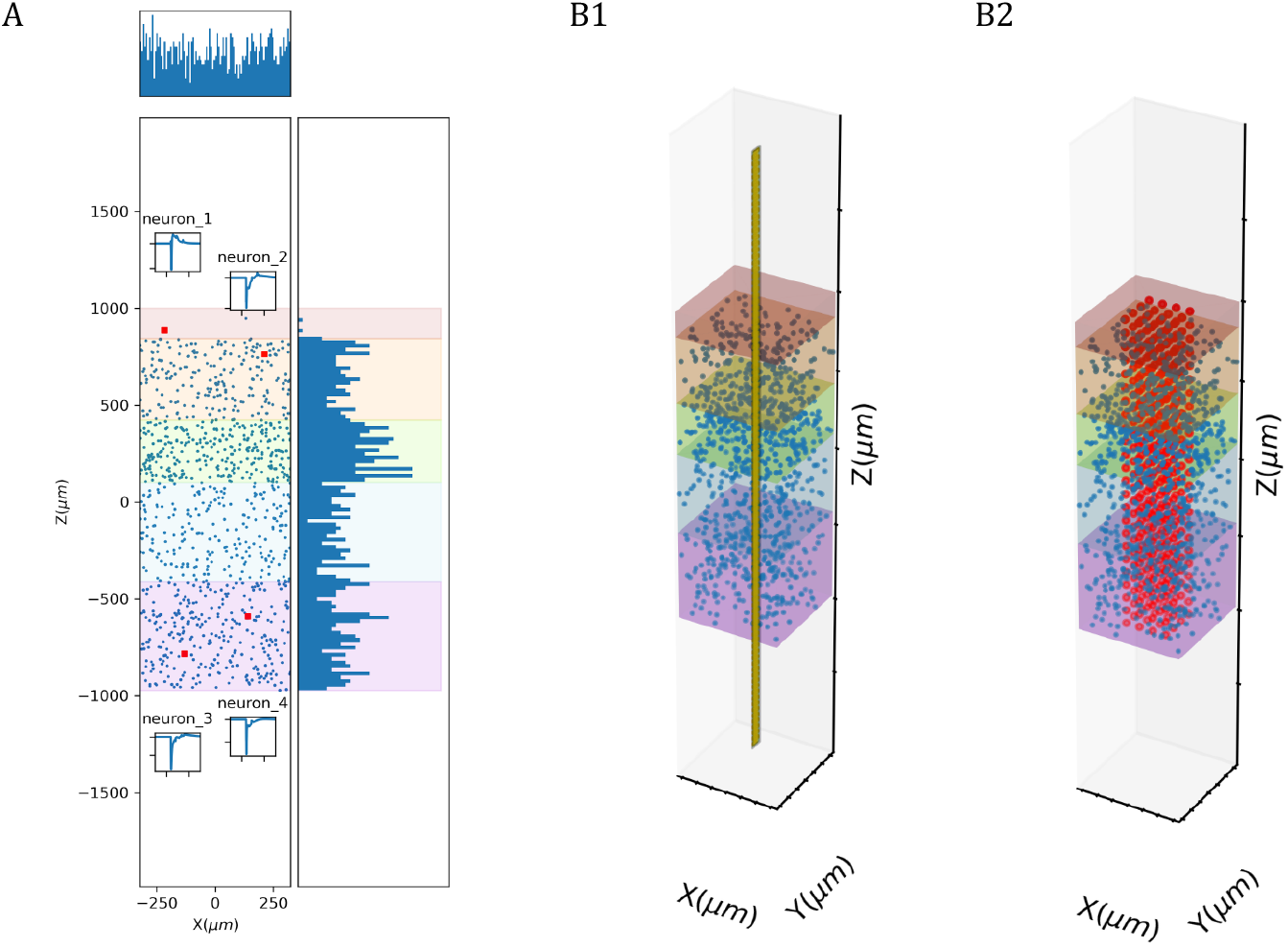
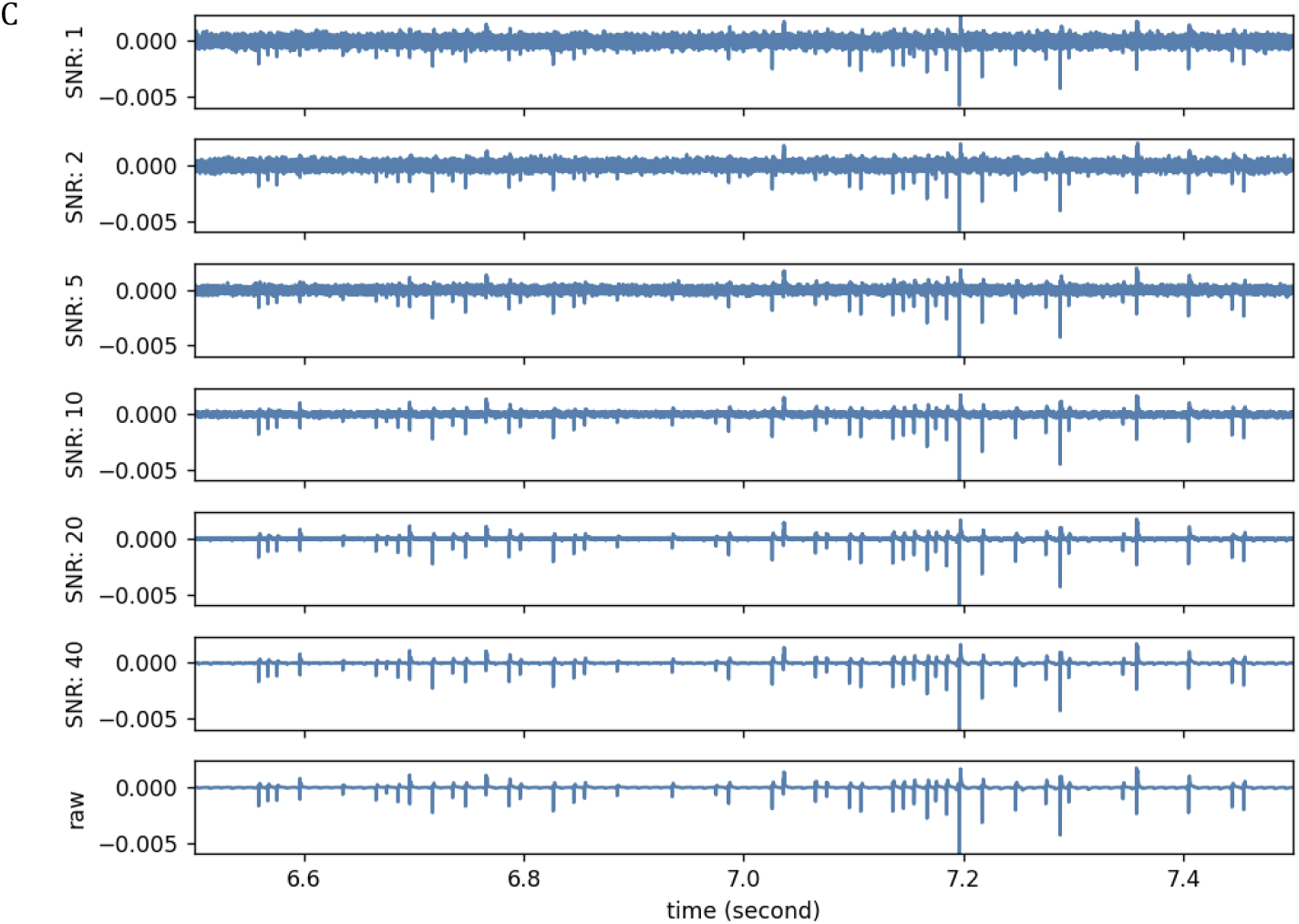
Simulated distribution of 1000 neurons network in a barrel column with 6 layers. (A) and 384-channel 2D linear electrode array (B1) and 384-channel 3D simple cubic electrode arrays (B2). (C) Increase the noise level by injecting white noise. From top to lower, signal-to-noise ratio of 1, 2, 5, 10, 20, 40, and non-injected raw data.

#### SNR of 40

We first looked into the best clean data, i.e. SNR of 40, the numbers of isolated units by Kilosort 2.0-3D (Fig. 4A). Since we know the ground truth, i.e. the locations of the 1000 neurons, we looked at the true-positive isolated units, meaning the units were designed and also isolated by Kilosort 2.0-3D. As expected, the performances of isolating units by 3D any electrode arrays were in general better (ca. 3 to 8 times higher) than NPX, and the performance was even better when the electrode pitch increased (from 45 μm to 150 μm) but saturated at ca. 125 μm. In particular, BCC and FCC isolated relatively more units than the other electrode arrays when the electrode pitch was 125 μm or 150 μm.

**Figure 4.**
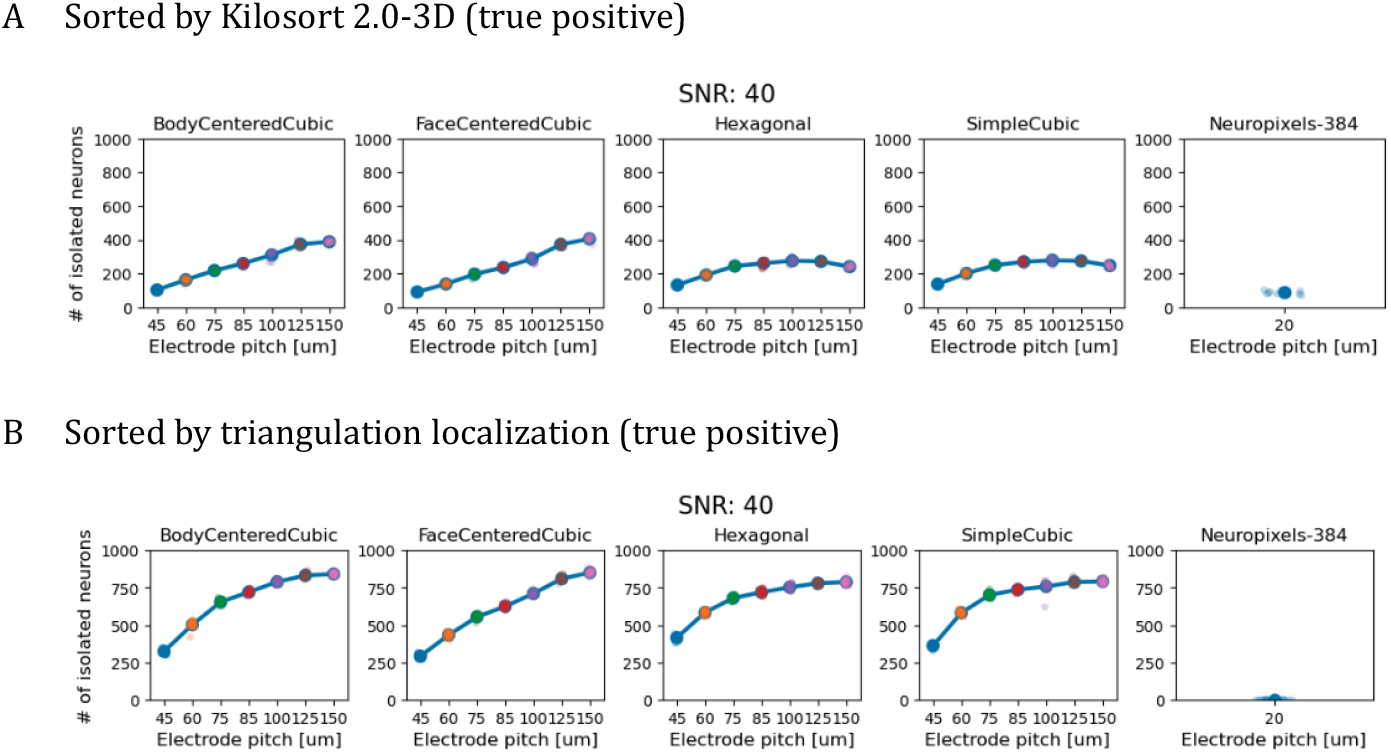
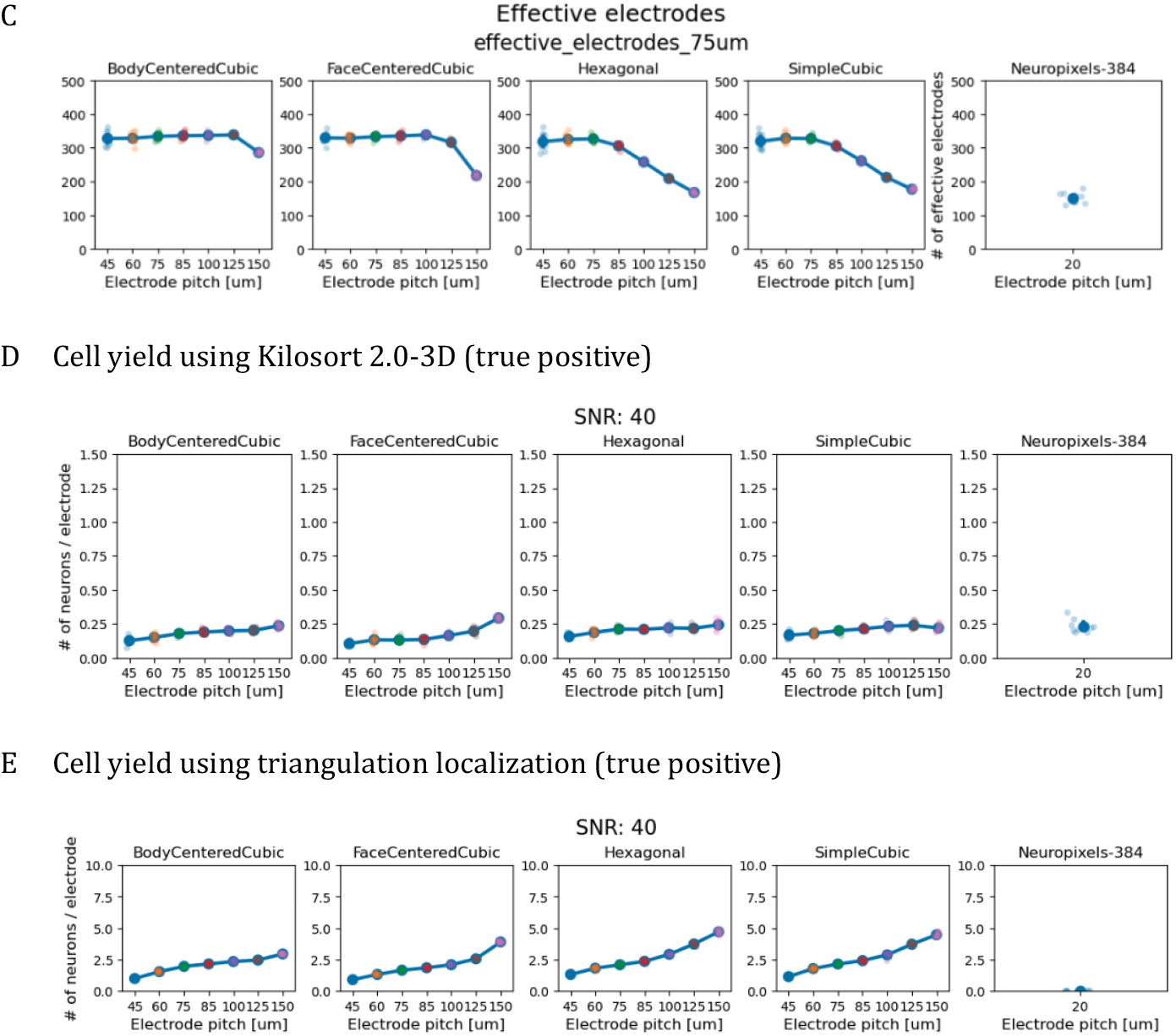
Benchmark of 4 types of 3D electrode arrays with electrode pitches of 45-150 um and Neuropixels probe in isolating neurons of simulated neural networks. 3D electrode arrays outperform NPX at an SNR of 40. (A) Using Kilosort2_3D for spike sorting, in general, all types of 3D electrode arrays outperform the Neuropixels probe, and isolate relatedly higher numbers of neurons with electrode pitches of 125-150 μm. (B) Using triangulation spike sorting, in general, all types of 3D electrode arrays outperform the Neuropixels probe. Note that the performance of isolating units using triangulation is even better when the electrode pitch is increased (from 45 μm to 150 μm), isolating up to 80% of neurons. (C) “effective electrodes” are defined as that electrodes are close to at least one of the 1000 neurons in 75 μm. As expected, the number of effective electrodes of 3D electrode arrays dropped from 330 to 180 (simple cubic) while the NPX has, on average, ca. 150 effective electrodes. (D) Cell yield is defined as the number of isolated units using Kilosort 2.0-3D (A) divided by the number of effective electrodes (C). (E) Cell yield is defined as the number of isolated units using triangulation localization (B) divided by the number of effective electrodes (C).

We then examined the true-positive isolated units by triangulation (Fig. 4B). Surprisingly, the performances of isolating units by 3D electrode arrays were much better (ca. 6 to 15 times higher) than NPX, and the performance was even better when the electrode pitch increased (from 45 μm to 150 μm) but saturated at ca. 125 μm. Similar to the results of using Kilosort 2.0-3D, BCC and FCC isolated relatively more units than the other electrode arrays when the electrode pitch was 125 μm or 150 μm.

However, it might be an unfair comparison since many electrodes of the NPX are far from the simulated neurons (Fig. 2B1). Thus, we defined “effective electrodes” where electrodes are close to at least one of the 1000 neurons in 75 μm. As expected, the number of effective electrodes of 3D electrode arrays dropped from 330 to 180 (simple cubic)-300 (BCC) while the NPX has, on average, 150 effective electrodes (Fig. 4C and Table S3). Next, we normalized the numbers of isolated neurons by Kilosort 2.0-3D and triangulation by the effective electrodes and defined them as cell yield (Fig. 4D and 4E). When using Kilosort 2.3D for spike sorting, the cell yield of 3D electrodes and the NPX are similar, at ca. 0.2. However, when using triangulation spike sorting, the cell yield of 3D neural probes increased from ca. 1 to 4 (i.e. FCC), respective to electrode pitch from 45 μm to 150 μm while the cell yield of NPX was worse, down to 0.1.

When we compared the ability of Kilosort2.0-3D and triangulation in isolating units, in the condition of 40 SNR, triangulation significantly isolated many more units per electrode. Thus, the simulation data strongly suggest that at 40 SNR, 3D neural probe (of FCC), together with triangulation, could isolate up to 80% of neurons and has a cell yield of up to 5, which is, to our knowledge, much higher than standard 2D electrodes (ca. 0.5).

#### SNR of 10

Next, we would decrease the SNR from 40 to 10 (closer to real ephys recording) and test its impact on the ability of the 3D neural probe and the NPX in isolating units. As expected, using Kilosort 2.0-3D, the number of isolated units dropped in 3D neural probes and the NPX (Fig. 5A), compared to SNR 40 (Fig. 4A). For example, the number of isolated units decreased from ca. 400 units to 300 units in FCC with electrode pitch of 150 μm. And, the cell yield also dropped from ca. 0.3 to 0.25 (ANOVA test, Fig. 5C).

**Figure 5.**
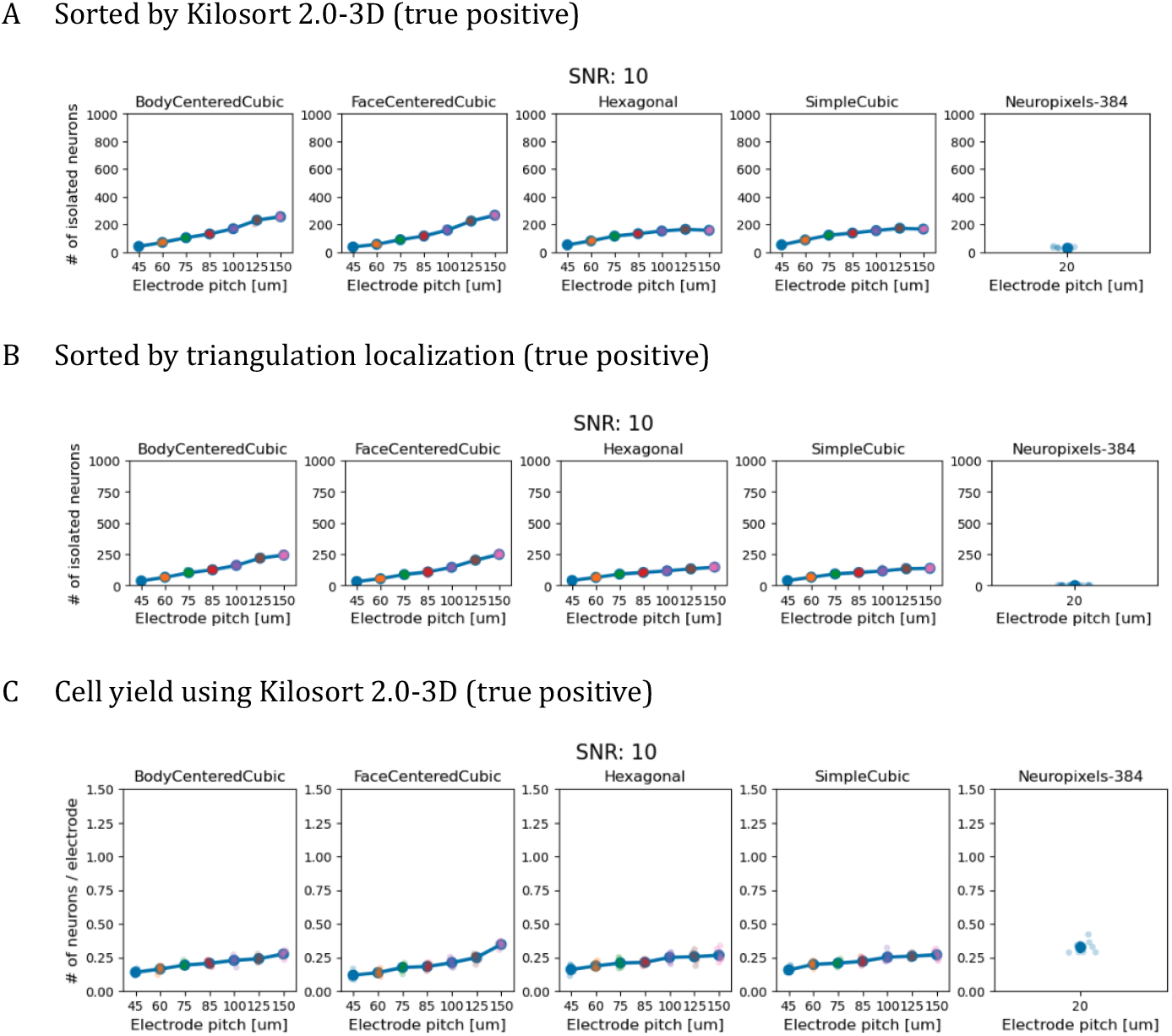
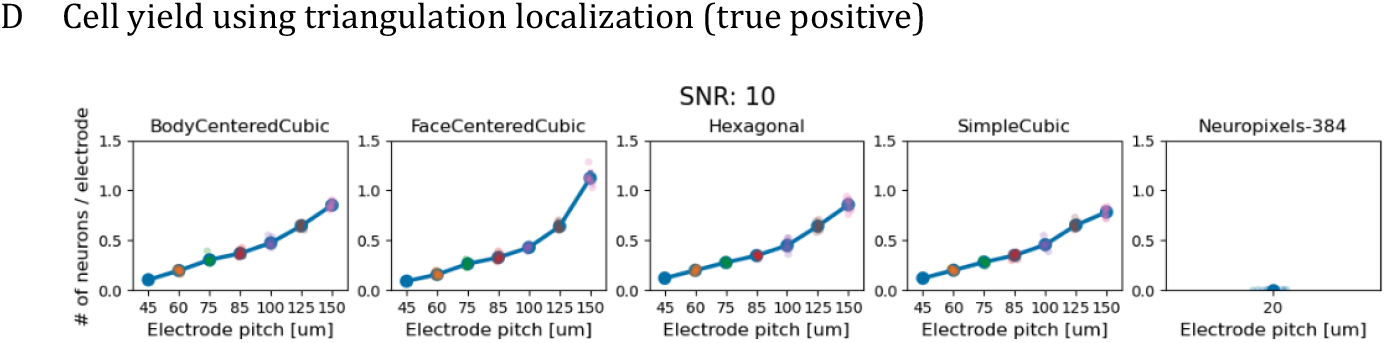
3D electrode arrays outperform NPX at an SNR of 10.

However, SNR seems to have a stronger impact on the triangulation spike sorting, leading to a decrease of isolated units from ca. 800 units (Fig. 4B) to 250 units (Fig. 5B). For cell yield, it dropped from ca. 4 to 1.1 (Fig. 5, FCC). Nevertheless, the spike sorting approach using triangulation for 3D neural probes seems to give a better yield than using Kilosort 2.0-3D. Similar to the SNR of 40, when using triangulation spike sorting, the cell yield of 3D neural probes increased from ca. 0.1 to 1.1 (i.e. FCC), respective to electrode pitch from 45 μm to 150 μm while the cell yield of NPX was worse, down to ca. 0.05.

#### SNR of 1

Next, we would decrease the SNR further from 10 to 1 (Fig. 6C, most upper panel), mimicking more noisy recordings, and test its impact on the ability of the 3D neural probe and the NPX in isolating units. As expected, using Kilosort 2.0-3D, the number of isolated units sightly dropped in 3D neural probes and the NPX (Fig. 6A), compared to SNR 10 (Fig. 5A). For example, the number of isolated units decreased from ca. 300 units to 200 units in FCC with electrode pitch of 150 μm. And, the cell yield also dropped from ca. 0.3 to 0.25 (ANOVA test, Fig. 6C).

**Figure 6.**
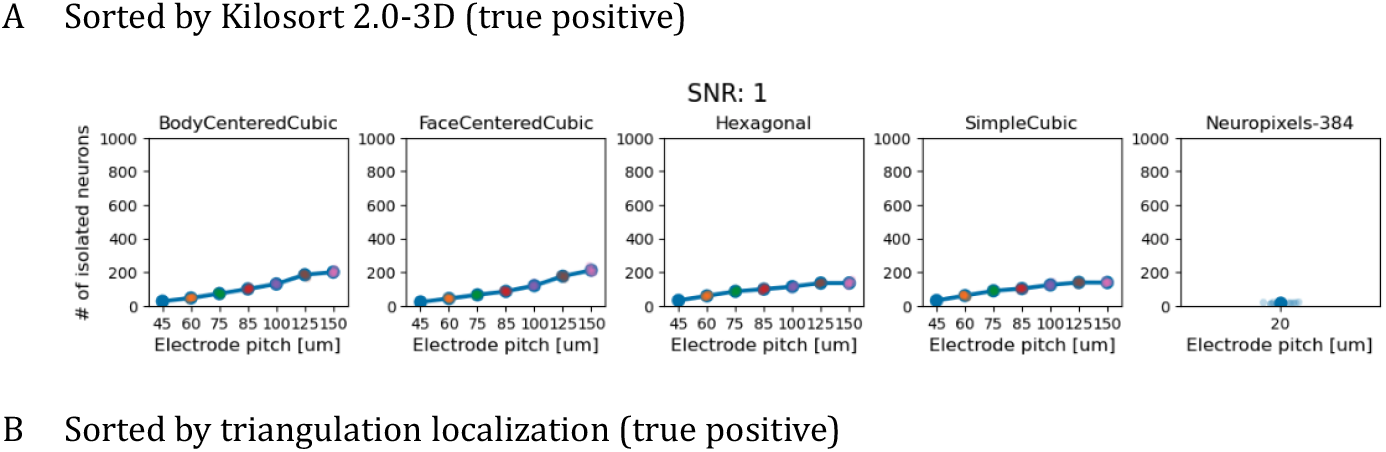
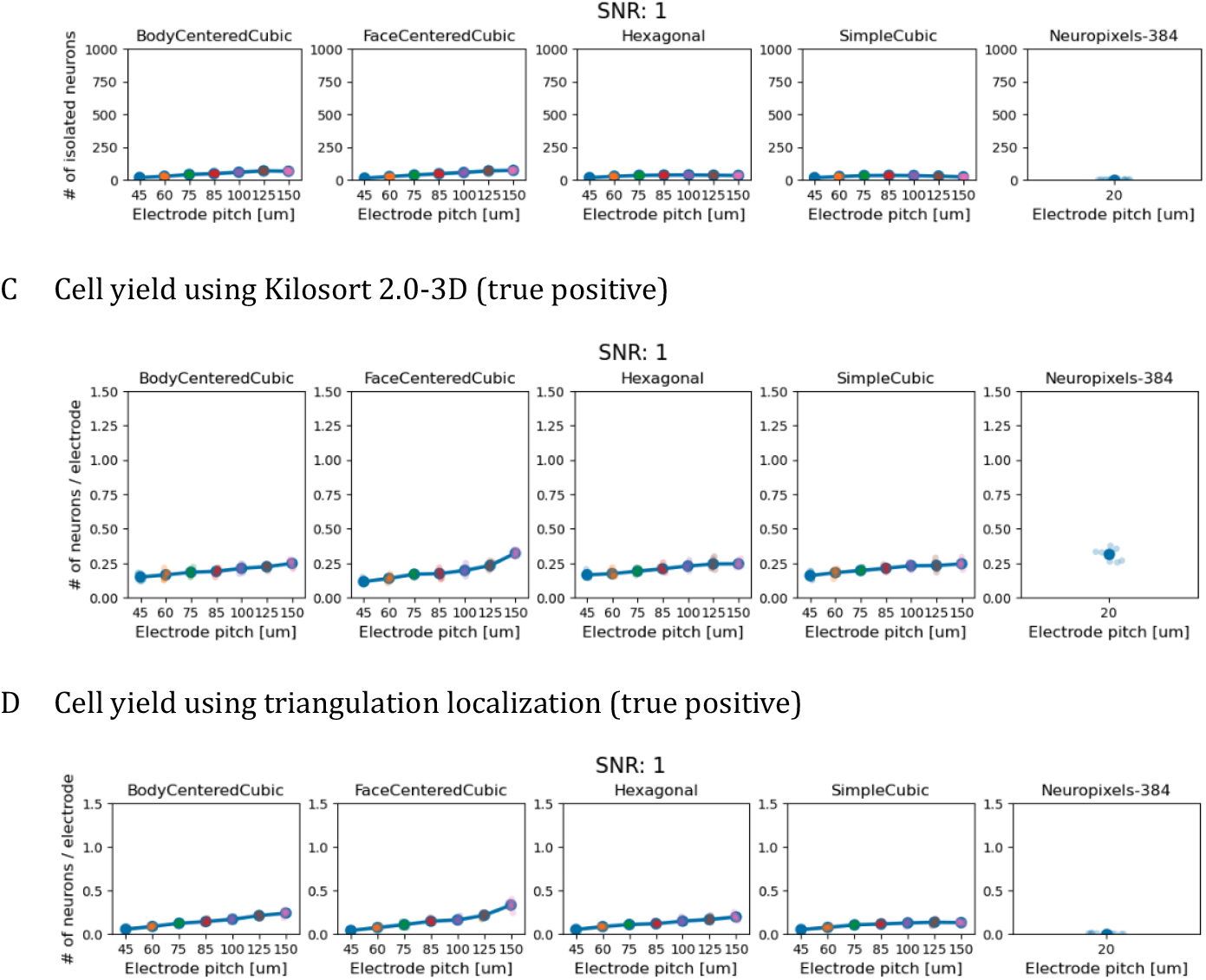
3D electrode arrays outperform NPX at an SNR of 1.

However, SNR seems to have a stronger impact on the triangulation spike sorting, leading to a decrease of isolated units from ca. 250 units (Fig. 5B) to 100 units (Fig. 6B). For cell yield, it dropped from ca. 1.1 to 0.4 (Fig. 6D, FCC). Thus, the spike sorting approach using triangulation for 3D neural probes seems to perform similarly to using Kilosort 2.0-3D for NPX (Fig. 6C).

### Ideally for real-time spike sorting

One more feature of triangulation localization is its speed. Since only the peak value of spikes are required for triangulation localization, the computing loading is much less than spike waveform-based spike sorting approach, e.g. Kilosort. We examined the time spent on triangulation localization, on average, it spent 80 ms to localize one unit, which is much fast than transitional spike sorting approach.

### Significant volume contributes to triangulation localization

Since the electrode pitches were highly correlated to the significant volume (Fig. 2, sensing radius of 150 μm; BCC and FCC) and the isolated units using triangulation (Fig. 4B; BCC and FCC), we, next, tested the hypothesis that the significant volume (Fig. 2) correlates to the results of triangulation localization (Figs. 4B, SNR: 40; 5B, SNR: 10; 6B, SNR: 1).

We summarized the following results.

1. These results of computing the significant volume of 3D electrode arrays suggest that steric probes provide one more degree of freedom than planar probes do, which can endow researchers with better spatial features for spike sorting, e.g. triangulation localization.
2. The simulation data strongly suggest that at a SNR of 40, 3D neural probe (of FCC), together with triangulation, could isolate up to 80% of neurons and has a cell yield of up to 5, which is, to our knowledge, much higher than standard 2D electrodes with Kilosort or triangulation.
3. With an SNR of 10, which is close to the real world, 3D electrodes with FCC arrangement may offer a better cell yield.
4. SNR, i.e. SNR of 1, has a stronger impact on the triangulation spike sorting, leading to a decrease in isolating units to the level of using the NPX and Kilosort 2.0-3D.
5. The significant volume is correlated with the ability of triangulation localization when the sensing radius was 150 μm.
6. Since only the peak values of spikes are required for triangulation localization, the computing loading is much less than the spike waveform-based spike sorting approach, e.g. Kilosort. Thus, 3D electrode array with triangulation localization is ideal for real-time spike sorting.

## Discussion

Following the summary of the results, we discuss the following topics.

### Any other study investigated the effect of electrode geometry on spike sorting performance?

Optimizing the design of electrodes on neural probes to improve cell yield involves various innovative approaches and materials. Several tools and techniques have been developed to enhance electrode performance and interaction with neural tissues. Conductive polymer coatings[34], and biomolecule integration[35]designs are key approaches that enhance electrode performance and interaction with neural tissues, leading to improved cell yield and recording stability. [36] is the first to systematically investigate the effect of probe geometry in 2D on spike sorting performance. However, to the best of our best knowledge, there is no bottom-up approach to design such an electrode array in 3D in order to have a higher cell yield.

### Discuss our neural network simulation and compare it to alternative tools

When simulating electrode recordings in neural networks, several tools beyond NEURON and LFPy are available. These tools offer diverse functionalities for modeling and simulating neural activity and electrode interactions. For example, the Virtual Electrode Recording Tool for EXtracellular potentials (VERTEX[37,38]) is a MATLAB-based tool that simulates LFPs from large neuron populations. It allows for virtual electrode placement in 3D and can simulate the effects of electrical stimulation on neural circuits.

### The exploration volume for Kilosort and the significant volume for triangulation localization?

Data in Fig. 8 showed a high correlation between significant volumes and performance of triangulation localization. However, the significant volume is not critical for Kilosort (data not shown), but the exploration volume is. Thus, in the aspect of simulations, for traditional 2D electrode arrays, exploration volume should be the better parameter when designing any type of 2D electrode array; the larger exploration volume, the better. However, for design 3D electrode arrays, the critical volume should be applied; the larger critical volume, the better.

### Discuss why triangulation spike sorting is for 3D probes while Kilosort, etc., is for 2D probes

Kilosort was originally designed to perform spike sorting for 2D electrode arrays. In this study, we rewrote it for 3D electrode array (as Kilosort 2.0-3D, see Fig. S1). However, its performance in spike sorting for 3D electrode arrays was worse than triangulation localization (see Fig. 5, SNR: 40). However, it is more resistant to background noise (see Figs. 5, SNR: 10; Fig. 6, SNR: 1). Therefore, when the background noise is strong, Kilosort seems a better tool in solating units than triangulation localization.

The triangulation location used in 2D electrode arrays is shown[33]. The authors claimed triangulation localization improved motion correction for Neuropixels recordings but did not investigate the cell yield. [39] fabricated a 3D silicon probe with 1024 contacts (Its x−y−z electrode pitch is 250−12−350 μm, and the volume enclosed by the array is 750−756−1050 μm) and demonstrate a cell yield of 2.5 using spike waveform-based spike sorting. Unfortunately, its electrode pitch is up to 350 μm, which is not ideal for triangulation localization. Thus, based on the assumption that each electrode can only detect spikes originating within a sphere of radius (sensing radius, up to 100 - 150 μm, we should design and fabricate 3D electrode arrays with electrode pitch all (x−y−z electrode pitch) within 150 μm.

### Why do 3D probes with triangulation localization outperform 2D probes in cell yield?

Triangulation localization largely replied on the critical volume, Figs. 1 and 7. Although triangulation localization can be performed for spikes recorded with 2D electrode arrays, but its significant volume is limited [33]. Adding one more dimension can significantly increase the significant volume. In this study, we demonstrated that FCC is a potential 3D structure for designing electrode arrays in 3D.

**Figure 7.**
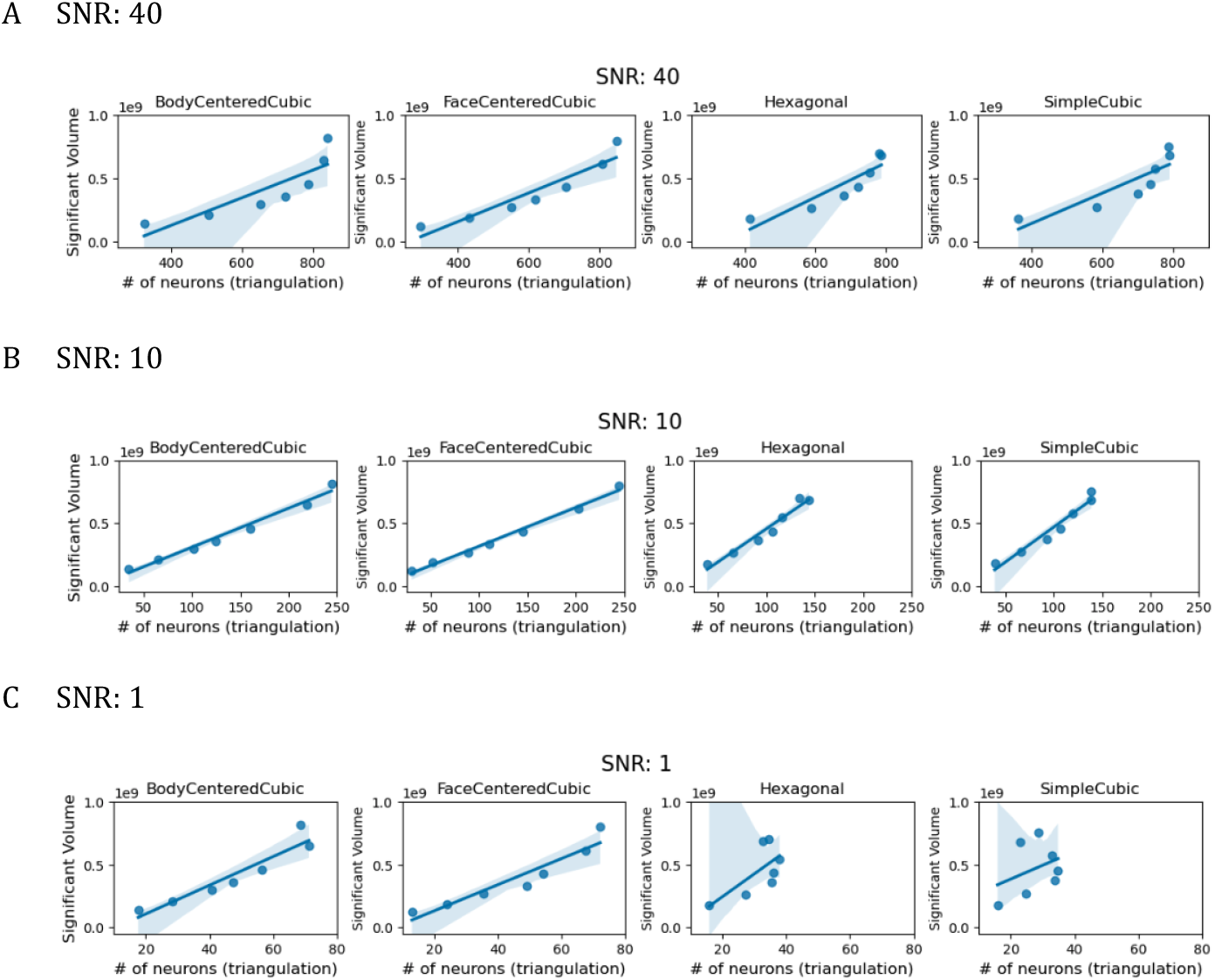
Correlation between significant volumes and performance of triangulation localization.

### Future challenges and prosepectives

1. Since the technology of making electrodes would be improved over time, one should extend electrode sensing distance up to 500 μm to calculate the significant volume.
2. Since the background noise has a strong impact on the triangulation localization, can the algorithm of triangulation localization be improved? How can we further reduce electrode impedance?
3. Real-time triangulation localization for high-channel count 3D electrode arrays might require much more computer power. Can we use GPUs to distribute the jobs for laege-scale real-time triangulation localization?
4. In this study, we simulated a neural network for a barrel column. However, if one targets the hippocampus, how should the neural network be adapted in a fast manner in order to opt for a better 3D neural probe?
5. How can such complex 3D electrode arrays be made?

## Methods

### Preparing of simulated neural signal

In-silico simulation reduces sacrifices of animals and fabrication time of prototype probes. Neuron[40] and LFPy[41] are adopted to generate virtual neural signals in this work. A cube of 646μm×646μm×3970μm with 1000 neuron cells in it is prepared to mimic neuronal activation in cortical column[42]. The neuron models are obtained from The Neocortical Microcircuit Collaboration Portal of Blue Brain Project[43]. These neurons are distributed uniformly in both x and y direction. Furthermore, these neurons were distributed among six layers in z direction which corresponded to six-layered Neocortex. The detailed configuration is summarized in Table1. To suppress the influence of signal collision, the firing interval of neurons is set as 5ms. In order to benchmark the performance of steric neural probes and traditional planar probes, the same simulation configuration is used in all kinds of electrodes. Figure X. illustrates the arrangement of neurons and the relative position between neurons and electrodes.

### Injection of AWGN

Computer simulations were employed to generate noise-free simulated signals. To facilitate an objective comparison between spike sorting and triangulation algorithms using actual experimental data, we injected Additive White Gaussian Noise (AWGN) into the simulated signals. The implementation of AWGN was based on the following formula

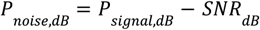

in which *P*_*noise,dB*_, *P*_*signal,dB*_ represent the average noise power and average signal power in dB, respectively. *SNR*_*dB*_ is the parameter enabling us to achieve varying signal-to-noise ratios (SNRs). Higher SNRs corresponded to lower levels of Gaussian Noise. We generated simulated signals with SNRs of 40 dB, 20 dB, 10 dB, 5 dB, 2 dB, 1 dB, and 0.1 dB, as illustrated in Figure 4(a)~(f). By doing so, we aimed to approximate real-world scenarios while concurrently evaluating the noise tolerance capabilities of spike sorting and triangulation algorithms. This approach allows us to gain insights into the efficacy of these techniques when confronted with different noise levels in practical experimental settings.

### Significant volume and exploration volume

We introduced a flag counter to determine the exploration volume and the overlapped explorable space of adjacent probes. We used feature flags to distinguish whether or not a voxel is located within the explorable area of a probe. The flag of a certain voxel will be plus 1 if the distance between the voxel and the center of the electrode is less than 75μm. The total amounts of voxels with non-zero flags represents the size of the exploration volume. The value of the flag counter represents the overlapping degree(OD)of the explorable space of adjacent probes. The higher OD value of a voxel means that more electrodes can detect the spike activity of one neuron located on the voxel.

### Kilosort2.0-3D

Kilosort is a computational tool for spike detection and sorting, widely used in neuroscience to analyze neural signals recorded from high-density electrode arrays. It was developed by Marius Pachitariu and his team at Cortexlab, University College London (UCL), and he is currently affiliated with HHMI Janelia. Utilizing template matching, it can classify spikes, even distinguishing overlapping waveforms. By GPU acceleration and dimensionality reduction of singular value decomposition (SVD), Kilosort significantly enhances the computational efficiency of large-scale data processing. Since the release of version 2.0 (October 2020), Kilosort has undergone continuous development to accommodate a broader range of signal types, leading to subsequent releases of versions 2.5 (January 2021), 3.0 (October 2022), and 4.0 (April 2024). With its automated spike sorting workflow, Kilosort minimizes the need for manual calibration, making it a powerful tool for large-scale neural data analysis.

Kilosort first preprocesses the recorded signals by organizing the data and performing mean-centering, followed by using filters to remove low-frequency noise and applying whitening to reduce common noise. After detecting spikes through thresholding, Kilosort applies singular value decomposition (SVD) to perform dimensionality reduction and clustering analysis on waveform fragments, thereby generating initial templates. Spike fragments are matched to these initial templates, with spikes reassigned iteratively to optimize the result. Finally, similar templates are merged, and minor templates are removed, yielding the final spike clustering result.

Kilosort is primarily designed for analyzing signals from planar electrodes, requiring two-dimensional information for input signals and channel coordinates. To enable its application to signals recorded by three-dimensional electrodes, we extended the spatial coordinate related calculations in the algorithm from two-dimensional to three-dimensional coordinates. This adaptation allows Kilosort to perform spike detection and sorting on three-dimensional datasets effectively.

#### Enhanced Kilosort from 2D to 3D

Since the Kilosort program primarily analyzes and processes data based on signal correlations and the relative distances between electrodes, we enhanced the version 2.0 program by extending its two-dimensional coordinate system, defined by X and Y coordinates, into a three-dimensional coordinate system with the addition of a Z coordinate. This modification enables the calculation of spatial relationships in three dimensions, allowing the program to process three-dimensional signals. To validate the accuracy of the modified three-dimensional version, we compared its computational results with those of the original version. Through testing with example datasets, we examined the clustering outcomes for signals.

### Triangulation spike sorting

It is shown mathematically that we only need four non-coplanar electrodes to determine the location of an excited neuron in three-dimensional space. In practical terms, the two intersections of three electrodes are very closed (less than 10μm) so that we can use only three electrodes to locate a neuron.

## Code availability

Our NEURON and LFPy implementation of neuron simulation is available by request.

## Acknowledgements

The authors would like to thank the National Center for High-performance Computing of Taiwan for providing computational and storage resources. This work was supported by the following grant: NSTC 113-2640-B-007-001, Taiwan.

## Notes

### Competing Interest Statement

The authors have declared no competing interest.

