## Supplementary material for "Real-time spike sorting with 3D neural probe and triangulation localization": Fig. S1

### Suppl. Figures

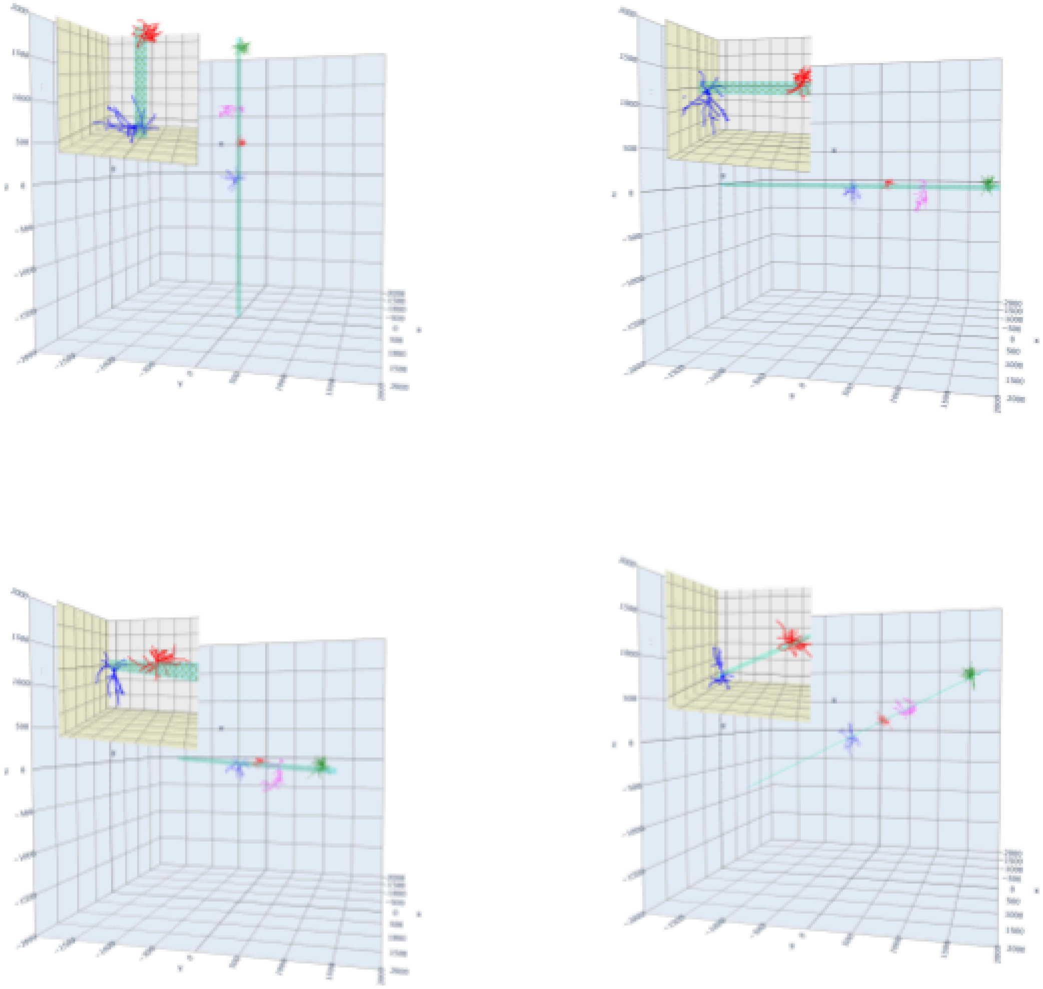

| | Electrode Locations | Number of well isolated Clusters |
|---|---|---|
| Kilosort 2.0 | 2-D | 15 |
| Kilosort 2.0 (Extend to 3-D) | 3-D | 15 |
| | 3-D (Rotation X1Y1Z1) | 15 |
| | 3-D (Rotation X2Y2Z2) | 15 |
| | 3-D (Rotation X3Y3Z3) | 15 |

Figure S1. Validation of Kilosort 2.0-3D. (A) Illustrations of orientation of a Neuropixels probe in 2D (a), 3D (b), 90-degree rotated to y-axis (c), further 45-degree rotating to z-axis (d), and further 45-degree rotating to x-axis (e). Inserts in each panel indicate magnification of electrodes at the tip of the Neuropixels probe. (B) After rotating the probe in different angles in space , Kilosort 2.0-3D is capable of isolating the same number of well isolated clusters, which validates the use of Kilosort 2.0-3D in 3D electrode arrays.
